# Biophysics of the Coronavirus—Membrane Interaction—Role of Nonequilibrium Binding Energy

**DOI:** 10.1101/2025.06.10.658990

**Authors:** Ikechukwu Iloh Udema

**Affiliations:** Department of Chemistry and Biochemistry, Research Division, Ude International Concepts Ltd., 862217, B. B. Agbor, Delta State, Nigeria

**Keywords:** SARS-CoV-2 variants, diffusivities, thermodynamic temperatures, viscosity, mucus, path function, cytoplasm, nonequilibrium binding energy

## Abstract

Scientists seemed not to have explored mechanical kinetic energy, a path function for the determination of nonequilibrium binding energy (NEBE). The study explored mechanical kinetic energy, a path function in particles that requires nonequilibrium binding energy (NEBE) to counteract it. It aimed to show that there was a minimally sufficient NEBE to counteract mechanical kinetic energy, using literature data to evaluate derived equations and computations. As is typical, diffusivities increase with temperature; however, near binding, the binding limitations cause the diffusivities (1.519 → 2.784 *exp*. (− 15) m^2^/s for D variant; 1.415→2.577 *exp*. (− 15) m^2^/s for G variant) to be lower than when they are far from binding (4.36 →7.99 *exp*. (− 15) m^2^/s for D variant; 4.151→7.611 *exp*. (− 15) m^2^/s for G variant). With breath emission equal to 1.29 *exp*. (7)/m^3^, the NEBEs were 656.020 and 663.212 kcal/mol for Delta and Omicron variants of SARS-CoV-2, respectively; and with 27.9 *exp*. (7)/m^3^, the corresponding values were 568.921 and 633 kcal/mol; with a breath emission rate equal to 9.31 exp. (+6)/hr., the NEBEs for the Omicron variant in 1 hr. and in 1 min. were 651.703 and 683.353 kcal/mol, respectively; with 201 *exp*. (+6)/hr., the corresponding values were 444.157 and 663.212 kcal/mol. The G variant of SARS-CoV-2 showed higher NEBEs (952→671 kcal/mol) than D variants (834→588 kcal/mol), corresponding to 293.15→318.15 K. The decreasing trend in maximum NEBE 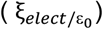 with rising temperature implies that the binding affinity of the virus may be attenuated at higher temperatures. It is, therefore, medically plausible to administer airborne, parenteral, oral, etc., drugs at temperatures above body temperatures that are tolerable and in a controlled fashion. Future studies should be directed to a definite determination of the size and molar mass of variants of SARS-CoV-2.

## 1. INTRODUCTION

It is well known that the speed at which any solute, be it either a drug or its metabolites or a pathogen like the recent SARS-CoV-2 that imprisoned humanity—which is feared, though it is not more deadly than Ebola—reaches its various targets should depend on the medium of transit. The medium offers resistance known as viscosity to motion driven by thermal energy. Beyond the retardative effect of viscosity are the compositional factors of the biological medium that further heighten the effect of viscosity. Since the model for nonequilibrium binding energy is linked to diffusivity, it is imperative to critique recent phenomenon bordering on diffusion and diffusivity of solute in a heterogeneous medium.

Referring to their unseen supplementary record (§3.1.), Marbach & Holmes–Cerfon (2020) were of the opinion that momentum relaxation (typically, loss of momentum due to obstacles, narrow inter-mucin fiber space, etc.) for micron-scale particles occurs over a timescale of *t*_*m*_ ≃ μs, while *on much longer timescales*, the equilibrium motion is diffusive with a diffusion coefficient independent of mass for large enough particles (Schmidt & Skinner, 2003 & Usabiaga et al., 2013). Inertia can only affect the short-time mobility of a particle (Usabiaga et al., 2013 & Bian et al., 2016). However, the dependency of the diffusion coefficient (or diffusion rate) on either mass or density is known to be inversely proportional to the square root and to the cube root of either the density or the mass of the particle for the small and large molecules, respectively; this is in line with Graham’s law of diffusion. It would appear, therefore, that the claim in the literature is conditional (this should not preclude new thinking).

Popovic (2022) explains that nonequilibrium thermodynamics links the rate of a biochemical process to its driving force, the Gibbs energy change. This bridges chemical kinetics and Gibbs free energy of binding. Mechanical kinetics, which plays a supporting role in catalyzed reactions, is not as well studied as (bio) chemical kinetics, which investigates rate constants for activation energies.

This study examined the mechanical kinetic energy required for virus or ligand binding on cell membranes. Meanwhile, a number of experimental techniques like fiber-optic-based fluorescence correlation spectroscopy, FB-FCS (Yamamoto & Sasaki, 2021); segmented fluorescence correlation spectroscopy, SFCS (Longo et al., 2024); single-particle tracking, SPT (Kapanidis et al., 2018); and fluorescence recovery after photobleaching, FRAP (Lorén et al., 2015), have been used to determine the diffusivity and corresponding viscosity known to reduce the former.

So far, experimental approaches have been explored for the determination of binding energy with its roots in electrostatic interaction energy, reported somehow as total energies (Nguyen et al., 2020). That the binding of the novel and old (SARS-CoV) viruses to ACE2 is driven by the electrostatic interaction (Nguyen et al., 2020) is not without evidence to the effect that at pH 7.4, used in an earlier work, the total charges of SARS-CoV-2-RBD and SARS-CoV-RBD are +3 and +2e, respectively (Nguyen et al., 2020). It has been demonstrated using steered molecular dynamics simulations (SMDSs) that a higher rupture force and extra pulling work are required to unbind SARS-CoV-2-RBD from ACE2-PD than to unbind SARS-CoV-RBD (Nguyen et al., 2020), similar to earlier findings (Wrapp et al., 2020). Regarding the equilibrium approach as a superior method for calculating absolute binding free energy (ABFE) implies a preference for it (Bhati et al., 2025). This is despite exploring mechanical process characteristics of a path function to determine binding free energy (Bhati et al., 2025). Despite externally forcing an equilibrated system out of equilibrium via nonequilibrium methods, such as nonequilibrium simulation (Serra et al., 2025), binding energy is reported as a kind of free energy. Scientists seem not to have explored mechanical kinetic energy, even if it is of electrostatic origin, a path function characteristic of any particle that must be opposed by a nonequilibrium binding energy. This study is predicated on the notion that state and path functions are separate.

The goal of this study was to show that, in contrast to the chemical kinetics-based energy that merely describes the thermodynamic stability or feasibility of fixing the ligand (such as viral spike protein) to the receptor (such as ACE2 in lipid rafts) before any further transformation, there is a minimally sufficient nonequilibrium binding energy to counteract the mechanical kinetic energy. To achieve the desired outcome, the following objectives were pursued: Using the derived equation, the theoretically optimal viral number density for infection, symptom outcome, and progression is determined; the viral number density in free or confined space and breath emission rate is mathematically related to multiplicity of infection (MOI) for a more practical or empirical determination of viral number density; the binding processes prior to infection are briefly described; the literature, with inference and therapeutic recommendations, regarding the impact of the medium’s composition and biophysical nature on viral infection is reviewed; and, finally, the nonequilibrium binding energy is computed.

## 2.0 BRIEF THEORETICAL BACKGROUND

### 2.1 Biophysical parameters-diffusivity, mean square displacement, and temperature

Given the molar mass of a solute, its translational diffusion coefficient at known temperature, the mean square displacement (MSD, (*ℓ*^2^)) (or the RMSD, *ℓ*) can be computed. Earlier computation (Udema, 2025) explored the following equation:

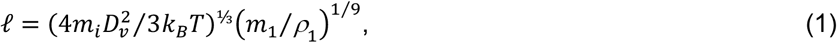

where *m*_*i*_, *D*_*v*_, *k*_*B*_, *T, m*_1_, and *ρ*_1_ are the mass of the solute (which can be viral particle), diffusivity, Boltzmann constant, thermodynamic temperature, mass of a molecule of water, and density of water respectively; (*m* /*ρ*)^1/9^ = *L*^⅓^. Solve for *D*_*v*_, to give:

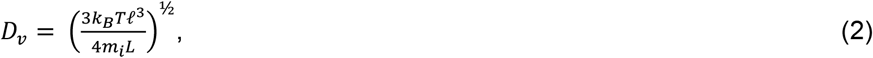

The time taken to cover such displacement is also computed with the following equation:

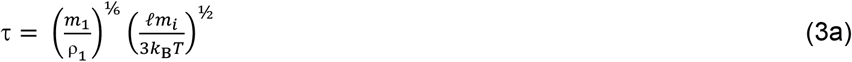

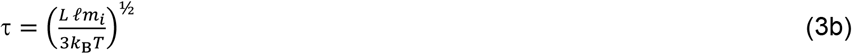

Equation is simplified to give:

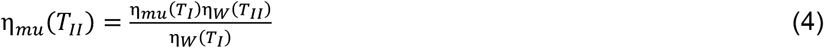

The approach in this study gives an insight into what may be expected in a practical experience. The advantage of the two equations is that information about the viscosity coefficient of the solution or media, such as cytoplasm, cell membrane, nucleoplasm, *etc*., is not required as long as the value of *D*_*v*_ is known. Yet the value of *D*_*v*_ should be for a known medium that has a verifiable viscosity.

The determination of the viscosity of the mucus at temperatures lower and higher than 25 ^0^C (293.15 k) is based on the equation as follows:

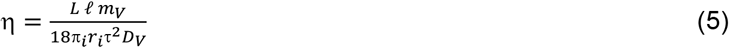

#### The viscosity coefficient based on Eqs (5) and (6)

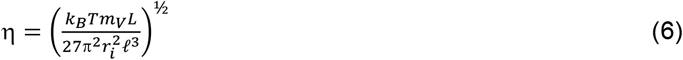

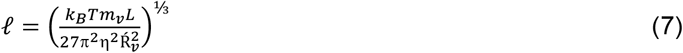

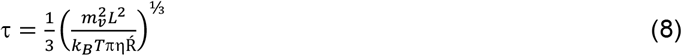

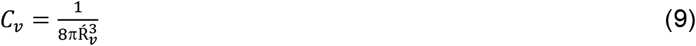

### 2.2 Viral number densities, short, and long intermolecular distances

where *C*_*v*_ and *Ŕ*_*v*_ are the theoretical maximum viral number density specific for a given virus and its hydrodynamic radius respectively.

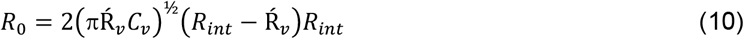

where *R*_0_ and *R*_*int*_ are the minimum interparticle distances wherein electrostatic attraction or repulsion commences and the maximum average intermolecular distance greater than *R*_0_.

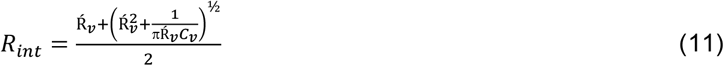

### 2.3 Small and large nonequilibrium binding energy

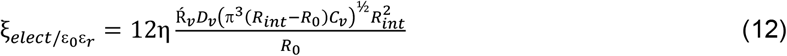

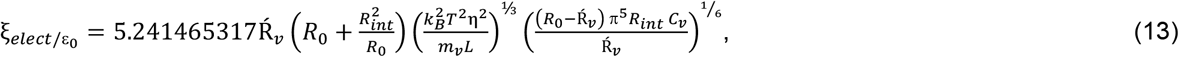

### 2.4 Multiplicity of infection, initial number of viral particles per cell, the corresponding viral number density

Leveraging on the idea that the multiplicity of infection (*MOI*) could be used to define the average number of virus particles that infect a cell (Shabram & Aguilar-Cordova, 2000), an equation derived in a manuscript under preparation is adopted in this study. This may obviate the misgiving of the term *MOI* by Shabram & Aguilar-Cordova (2000).

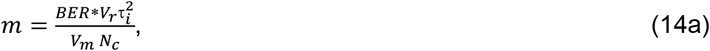

where *BER, V*_*r*_, *τ*_*i*_, *V*_*m*_, *N*_*c*_, and *m* are the breath emission rate per hour, ventilation rate per minute, period of time (taken to be one minute and one hour, for the purpose of this study) under consideration, molar gas volume, number of nasal cells, and multiplicity of infection (*MOI*). If on the other hand, the viral density (ℬ) in viral copies per cubic meter in the surrounding is known, the equation of *m* is given as:

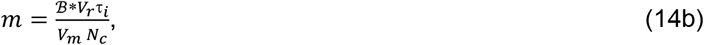

In this case the viral number density (*C*_*v*→ℬ_) per cell is given as:

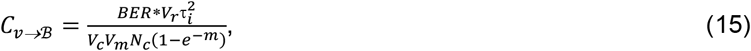

where *V*_*c*_ is the arbitrarily chosen volume of a cell as to imply that such, may be < or > the average volume of a mammalian cell. If the number density (or ℬ) in the air (which may be in the room, toilet, concert hall, *etc*.) rather than (*BER*) is given, then, the number density (*C*_*v*→ℬ_) of the virus per cell is given as:

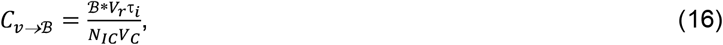

where *N*_*IC*_ = *N*_*c*_(1 − *e*^−*m*^). All these with detailed basis are again in a text book (Udema, 2025) in preparation.

Exploring different periods of time (one hour (a); one minute (b)) generates different values of *N*_*IC*_ and the corresponding *C*_*v*→ℬ_ values, initial interparticle distance (*R*_*int*(ℬ)_), initial interparticle distance (*R*_0(ℬ)_) wherein mutual attraction or repulsion may not occur because the number density is very low, narrower interparticle distance (*R*_*int*(*τ*→∞)_), and interparticle distance (*R*_0(*τ*→∞)_) wherein mutual attraction or repulsion may occur because the number density is very high.

### 2.5 Viral number density (*C*_*v*(*τ*→∞)_) as time tends to infinity given known number density of viral copies or breathe emission rate in the surrounding

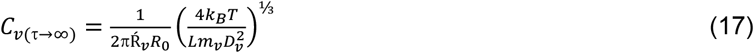

where *L* and *D*_*v*_ are the average intermolecular distance in the molar volume of water and the viscosity-dependent diffusivity at ambient temperature respectively.

### 2.6 Proposition that compositional factor determine diffusivity and ultimately, nonequilibrium binding energy, inferences, and therapeutic significance

Cross-linked, bundled, and entangled mucin fibers, which are released by goblet cells and the seromucinous glands of the lamina propria at the apical epithelium, are characteristics that can increase the viscosity of the mucus (Carlstedt & Sheehan,1989; Thornton & Sheehan, 2004). The mucin fibers, 3– 10 nm in diameter (Shogren et al., 1989), are proteins glycosylated via proline, threonine, and/or serine residues by O-linked N-acetyl galactosamine as well as N-linked sulfate-bearing glycans (Carlstedt & Sheehan, 1984). Glycan coverage of mucin is dense, with 25–30 carbohydrate chains per 100 amino acid residues (Lamblin et al., 1991), and make up the larger part of the dry mass (Masson & Heremans, 1973). Most mucin glycoproteins have a high sialic acid and sulfate content, which leads to a strongly negative surface that increases the rigidity of the polymer via charge repulsion (Shogren et al., 1989).

Proteins, specifically mucin fibres, increase mucus’s viscous and elastic moduli (Harbitz et al., 1984, Girod et al., 1992). This is supported by reduced transport rates for IgM and small IgA aggregates in mucus, slowed by low-affinity bonds with mucin (Saltzman, 1994, Olmsted et al., 2001). This suggests that SARS-CoV-2 mobility in mucus must be several-fold lower than in aqueous solution or pure water. Engineered polymeric nanoparticles with 200 and 500 nm diameters exhibited a 6-fold and 4-fold decrease in effective diffusivities compared to water, suggesting that larger molecular mass particles, like SARS-CoV-2, may show lower diffusivity in crowded viscoelastic environments (Lai, et al., 2007).

Amine-modified particles have been observed to undergo more rapid transport in cystic fibrosis sputum than carboxylated particles, an effect that may be caused by the reduced adhesivity of the neutrally charged amine–modified particles. The significance of this finding is that drugs, directly or indirectly as LNP–encapsulated drug can be made more mobile by being amine–modified so that the pathogens can be exposed to higher frequency of collision. “Agents that bear positive charge (amine-modified) may enhance viral mobility on ward to the cell membrane (this may be due to reduction of viral interaction with mucus fibers (Dawson et al. 2003) while those bearing negative charge may sink the virus into the mucus and retard their mobility towards the cell membrane”. Secondary bonds that form between interacting particles and mucus affect mostly the short time-scale transport of carboxylated particles, not the long-time scale transport, which was mostly diffusional ⟨Δ*r*^2^⟩/2*D*_*v*_∼ Δ*t* at high, albeit with a much diminished diffusion coefficient compared with diffusion in buffer (Norris & Sinko, 1997).

As noted in the literature (Dawson et al., 2003) the average particle transport rates of 100-, 200-, and 500-nm particles decreased with increasing time, a small but significant fraction of the smaller particles (100 and 200 nm) exhibited more diffusive transport rates *i*.*e*. (⟨Δ*r*^2^(Δt)⟩) ∼ Δt; *this seems to be an error, otherwise it is appropriate to state that* ⟨Δ*r*^2^⟩/2*D*_*v*_∼ Δt *where D*_*v*_ *and* ⟨Δ*r*^2^⟩ are the diffusivity and mean square displacement respectively). At long time scales (Δt >1 s), the ensemble average MSD of 100-and 200-nm particles scales with time, ⟨Δ*r*^2^⟩/2*D*_*v*_∼ Δt suggesting that particles eventually escape their cages and move more freely (Dawson et al., 2003). This study focuses on initial and final diffusivity values, similar to the state function, end states of diffusion, and binding period to achieve a non-state function. The study found that particles move in pores too small for 500-nm particles, supporting the claim that supramolecules like SARS-CoV-2 can only show diffusivity in mucus several times less than in water (Dawson et al., 2003). Examples of this are the G and D versions of SARS-CoV-2, which have, respectively, viscosity values equal to 2.5 and 2.7 exp. (−15) m^2^/s (Al-kuraishy *et al*. 2022). According to the literature cited thus far, medium heterogeneity—with varying microviscosities at different points—is responsible for subdiffusive events in SARS-CoV-2 variants.

### 2.7 Binding processes and cleavage of the S protein S1–S2 boundary by furin

The transmembrane S2 domain of SARS-CoV-2 mediates the fusion of viral and cellular membranes, with a polybasic cleavage site (PRRAR) required for efficient proteolytic cleavage. This site is essential for cell-cell fusion and viral entry into human lung cells. The infection begins with the binding of the spike protein to receptor-angiotensin-converting enzyme 2 (ACE2), resulting in viral RNA genome release (Hoffman et al. 2020, Ou et al., 2020, Walls et al., 2020, Xia et al., 2020, Lie et al., 2023).

## 3.0 EXPERIMENTAL

All of the equations used in this section were taken from the literature (Udema, 2025). Furthermore, since the arithmetic means (where appropriate) of the data in the literature were explored, there was no need for materials and equipment. Given different volumes of cells in the literature, a representative nasal mucus cell volume (similar to a median value) used in this study is 1125 μm^3^. Type I cells which are large squamous cells have a volume of ∼ 2,000 to 3,000 μm^3^ while Type II cells which are cuboidal cells have a volume of ∼ 450 to 900 μm^3^ (Crandall & Matthay, *et al*., 2001). The mean volume of the epithelial cells equal to 1256 +/- 240 (Uribe & Gunderson, 1997), 630 +/- 180 (for 15 different tissues) and 2330+/- 650 μm^3^ (for 12 types of isolated epithelial cells) (Devany *et al*., 2023) are known. The number of cells in nasal mucosa explored is exp. (9) (Crapo et al., 1982, Stone et al. 1992). The root mean square displacements (RMSDs) of the SARS-CoV-2 S protein explored at 0, 20, 40, and 60°C were 1.53, 2.51, 3.26, and 2.23 nm, respectively (Khan, *et al*., 2022). These are molecular dynamics simulation-based (MDS-based) results.

### 3.1 Methods

Since experiments were not conducted arithmetic means or otherwise of values in the literature were explored in this study.

#### 3.1.1 Testing the proposition that RMSD or MSD cannot be arbitrarily chosen

Equations (2), (3b), (7), and (8) were explored to test the proposition that RMSD or MSD cannot be arbitrarily chosen.

#### 3.1.2 Viscosities at temperatures lower and higher than 25 ^0^C (298.15 k)

Equation (4) was explored for the theoretical determination of viscosities at temperatures lower and higher than 298.15 K.

### 3.1.3 Multiplicity of infection

When the breath emission rate in viral copies per hour and the number density in viral copies per cubic meter in the surrounding air, whether in a room, outside, etc., are known, the multiplicity of infection (MOI, *m*) is computed using Eqs (14a) and (14b), respectively.

#### 3.1.4 The viscosity coefficient

The viscosity of the medium in which the diffusivity of any solute at a specified thermodynamic temperature is known or given is computed using Eqs. (5) and (6).

#### 3.1.5 Longer and shorter average interparticle distances wherein mutual attraction is either unlikely or likely

Longer and shorter average interparticle distances wherein mutual attraction is either unlikely or likely at low and high viral number density, respectively, are determined using the same model equation, such as Eq. (10).

#### 3.1.6 Maximum interparticle distance given specific viral number density

The maximum interparticle distance given any viral number density is computed using Eq. (11).

#### 3.1.7 Viral number densities per cell

##### 3.1.7.1 Optimum viral number density per cell

The viral number density that can give the shortest minimum interparticle distance for achieving a high-frequency collision of viral particles with the target cell membrane is computed using Eq. (9).

##### 3.1.7.2 Low viral number density per cell given BER and the number of viral copies per cubic meter in the surrounding

The low viral density, given the *BER* and the number of viral copies per cubic meter in the surrounding confined or open area, is computed based on Eqs (15) and (16), respectively.

##### 3.1.7.3 Higher viral number density given BER and the number of viral copies per cubic meter in the surrounding

Given either *BER* or viral copies per cubic meter in the surrounding, confined, or open environment, the higher viral number density per cell is calculated using Eq. (17).

### 3.2 Determination of the nonequilibrium binding energy with and without the influence of aqueous relative permittivity

Equations (12) and (13), respectively, were explored for the computation of the nonequilibrium binding energy before and after the viral particle’s effective collision with the membrane following the release of hydration water.

## 4.0 RESULTS AND DISCUUSION

This section begins with the report given in the literature. The rate at which cytoplasmic proteins and other solutes, as mentioned above, reach their various locations is usually limited by passive translational diffusion and the related diffusivity (Bellotto et al., 2022). Given any arbitrarily or intentionally selected time, bits of diffusivity information can improve the calculation of root mean square displacement (RMSD); in contrast to time, RMSD cannot be selected arbitrarily.

The importance of drugs is best appreciated, first, from the perspective of pharmacokinetics, which is primarily dependent on diffusivity, without which the intended effect of such drugs cannot be achieved; second, from the perspective of pharmacodynamics, which can be unrealizable if the drug, directly or indirectly through its metabolite, does not bind to the target object—cell membrane, protein, receptor, etc. It appears, therefore, that diffusivity is more or less “a bridge” between pharmacokinetics and pharmacodynamics; destroy the bridge and be faced with no distribution and binding to target macromolecules, cells, tissues, etc.

Models or equations can only become useful after their confirmation following a series of evaluations to verify their consistency and possible generalizability. In order to achieve this, RMSD (*ℓ*) and the time (*τ*) spent traveling such a displacement were calculated using Eqs. (1) and (3a/3b), respectively. Let’s begin: Using FRAP, the diffusion coefficient of bovine serum albumin in normal tissue grown in a thin transparent window in the ear of a rabbit had been determined (this is an example of direct experimental determination). The average albumin diffusion coefficients (in normal tissue), molar mass, and Stokes– Einstein radius were 5.8 +/- 1.3 *exp*. (−11) m^2^/s, 67 kg/mol, and 3.55 nm respectively(Chary & Jain, 1989). Assuming a Kelvin temperature of 310.15 K and substituting it into Eq. (1) to obtain:

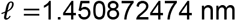

In order to determine the value of the average time taken to travel the computed RMSD, *ℓ*, the value above is substituted into Eq. (3a/3b) to give:

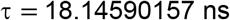

Next these values should be able to reproduce the literature (Chary & Jain, 1989) value of diffusion coefficient if the original Einstein equation (*D*_*i*_ = *ℓ*^2^/2τ) is fitted to the data to give:

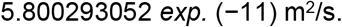

The 2021 CODATA (Tiesinga et al., 2021) values of Boltzmann constant and Avogadro’s number were explored. If the common values such as 6.02252 *exp*. (23)/mol. and 1.380485245 *exp*. (− 23) J/mol./K for Avogadro’s number and Boltzmann constant respectively, were explored the value of *D*_i_ is equal to 5.799999997 *exp*. (−11) m^2^/s. In all these computations, the viscosity coefficient was not explored. This important parameter is determined from two equivalent equations as follows: First, the viscosity coefficient is solved in both Eqs (5) and (6) as follows:

From Eq. (5):

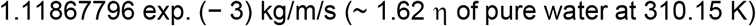

From Eq. (6):

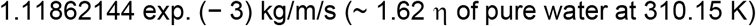

Both results show that the tissue (the type is not specified in the literature or probably not seen) is more viscous than pure water.

With nothing else except RMSD, the diffusivity and time expended travelling the displacement are shown in Table 1a. Given the RMSD, the diffusivity and time expended travelling the displacement were computed based on Eqs (2) and (3b) respectively. The values ranged from 3.633 to 12.099 *exp*. (−14) m^2^/s, with the peak value at 313.15 K following similar trend for RMSD reported elsewhere (Khans et al., 2022). The time spent traveling the RMSD ranged from 32.215 to 43.919 μs. The corresponding viscosity coefficients ranged from 0.54650284 (the lowest at 313.15 k) to 1.587349262 kg/m/s (the highest at 273.15 k).

**Table 1a.** Diffusivity and time of accomplishing predetermined RMSD (*ℓ*) at different temperatures: A test of non-arbitrariness of RMSD.

| T/K | 273.15 | 293.15 | 313.15 | 333.15 |
| --- | --- | --- | --- | --- |
| *$\ell$<br>(nm) | 1.53 | 2.51 | 3.26 | 2.23 |
| $D_v$ (Eq. (2))<br>(e. (-14) m<sup>2</sup>/s) | 3.633 | 7.909 | 12.099 | 7.060 |
| $\tau$ (Eq. (3b))<br>(μs) | 32.215 | 39.830 | 43.919 | 35.218 |
| $\ell^2/2\tau$<br>(e. (-14) m<sup>2</sup>/s) | 3.633 | 7.909 | 12.099 | 7.060 |
| $\eta$ (Eq. (5))<br>(kg/m/s) | 1.583 | 0.781 | 0.547 | 1.002 |
| $\eta$ (Eq. (6))<br>(kg/m/s) | 1.583 | 0.781 | 0.547 | 1.002 |
| RMSD (Eq. (7))<br>(nm) | 1.53 | 2.51 | 3.26 | 2.23 |
| $\tau$ (Eq. (8))<br>(μs) | 32.215 | 39.830 | 43.919 | 35.218 |
| $\ell^2/2\tau$<br>(e. (-14) m<sup>2</sup>/s) | 3.633 | 7.909 | 12.099 | 7.060 |

\* Represents data adopted from the literature (Khans et al., 2022); e means exponent (exp.)

Cholesterol affects the initial lipid phase, causing a decrease in viscosity in gel phases due to gradual gel phase transition into liquid-ordered (Lo) phase. This results in a rigid stem cell plasma membrane during differentiation (Wu et al. 2013, Dent et al., 2015, Meleshina et al., 2018, Matsuzaki et al., 2018, Kashirina et al., 2020). This may be applicable to other kind of cells. The RMSD and the corresponding time may be determined using, respectively, Eqs (7) and (8) given that the membrane and cytoplasm have viscosities of 416 cP (Shimolina et al., 2021) and 50 cP (Khismatullin et al. 2012), respectively. Thus as Table 1b, shows, the media presenting different viscosities expectedly showed root mean square displacement (RMSD), duration (*τ*) of traveling RMSD, diffusivity (*D*), translational velocity, terminal velocity, and the driving force, the thermochemical potential gradient that are different; the latter implies that chemical potential gradient is not enough to drive motion even at very low temperatures in which thermal energy is very low. There is a need to note inconsistences in the viscosities in the literature. Notable examples include stem cells billed for different tissues, such as osteogenic and chondrogenic tissues; Kashirina et al. (2020) reported, at day 14, 458.2 ± 37.02 cP and 438.04 ± 36.67 cP vs. 510.47 ± 40.27 cP in control for osteogenic and chondrogenic MSCs, respectively, given that advanced fetuses (or neonates) are also infested by SARS-CoV-2 or any other virus, even though adults, particularly the elderly, are most susceptible to viral infection. The drug, oxaliplatin, changes the viscosity of the plasma membrane of cancer cells that have been grown (Shimolina et al., 2021). If generalizable (or with any other formulated drug), it should be useful in controlling membrane viscosity. Other examples of different viscosities are 36.3 +/− 11.2 Pa.s (η_MCF-10A_); 65.9 +/−11.4 Pa.s (η_MCF-7_); 12.0 +/− 57 Pa.s (η_MDA-MB-231_) (Dessard et al., 2024) for different cell lines and 1-50 cP (Molines et al., 2022) probably at 22 ^0^C were also reported.

**Table 1b.**
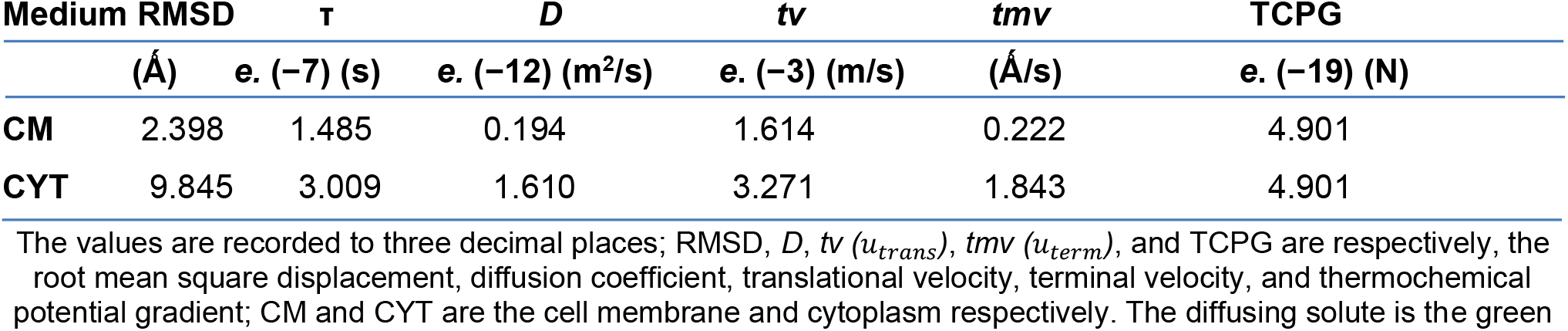
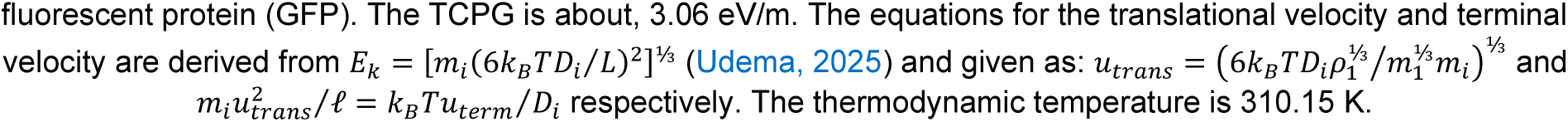
Biophysical parameters associated with CM and CYT.

| Medium | RMSD | $\tau$ | $D$ | $tv$ | $tmv$ | TCPG |
| --- | --- | --- | --- | --- | --- | --- |
| | ($\text{\AA}$) | e. (-7) (s) | e. (-12) (m<sup>2</sup>/s) | e. (-3) (m/s) | ($\text{\AA}$/s) | e. (-19) (N) |
| CM | 2.398 | 1.485 | 0.194 | 1.614 | 0.222 | 4.901 |
| CYT | 9.845 | 3.009 | 1.610 | 3.271 | 1.843 | 4.901 |

The values are recorded to three decimal places; RMSD, $D$ , $tv$ ( $u_{trans}$ ), $tmv$ ( $u_{term}$ ), and TCPG are respectively, the root mean square displacement, diffusion coefficient, translational velocity, terminal velocity, and thermochemical potential gradient; CM and CYT are the cell membrane and cytoplasm respectively. The diffusing solute is the green
fluorescent protein (GFP). The TCPG is about, 3.06 eV/m. The equations for the translational velocity and terminal velocity are derived from $E_k = [m_i(6k_BTD_i/L)^2]^{1/2}$ (Udema, 2025) and given as: $u_{trans} = (6k_BTD_i\rho_1^{1/3}/m_1^{1/3}m_i)^{1/2}$ and $m_i u_{trans}^2/\ell = k_B T u_{term}/D_i$ respectively. The thermodynamic temperature is 310.15 K.

SARS-CoV-2 begins its attractive interaction with the target cell membrane within the interparticle distance (*R*_0_) shown in Table 2. Table 2 also shows the average interparticle distance (*R*_*int*_) outside the region of mutual electrostatic effect for the D- and G-variants of SARS-CoV-2, as well as the ideal numerical density of the virus that is adequate (as opposed to the “dilute state”) to cause frequent collision with cell membranes.

**Table 2.** Physical parameters, interparticle distance and viral number density pertinent to cell-virus interaction.

| Average distance ($R_{int}$) between SARS-CoV-2 and target cell membrane | |
| --- | --- |
| SARS-CoV-2 (D-variant): | 101.054 nm |
| SARS-CoV-2 (G-variant): | ~ 109.138 nm |
| Optimum viral number density ($C_v$) | |
| SARS-CoV-2 (D-variant): | ~ 3.083 exp. (+20)/m<sup>3</sup> |
| SARS-CoV-2 (G-variant): | ~2.448 exp. (+20)/m<sup>3</sup> |
| Interparticle distance ($R_0$) wherein mutual attraction begins | |
| SARS-CoV-2 (D-variant): | ~71.456 nm |
| SARS-CoV-2 (G-variant): | ~ 77.172 nm |

\* (the data for SARS-CoV-2 D and G variants are from the works of Al-kuraishy *et al.* (2022))

A common observation, as seen in Table 3, is that as the Kelvin temperature rises, the diffusivities rise but the viscosities fall. The mobility of both beneficial and detrimental biomolecules is affected by this. For example, germs are more likely to move through a medium like nasal mucus that has a higher viscosity than water as the temperature rises. Although cholesterol plays a dual role in stiffening the cell membrane (CM) at melting temperatures and fluidizing it at lower temperatures, the platform for the virus’s binding is likely to be thermodynamically impossible when the CM is reached at temperatures higher than body temperature. Regardless of the magnitude of viscosity, increasing temperature alters the prevailing diffusivities. There is always an increase in diffusivities with increasing temperature. This is what Tables (3) and (4) illustrate.

**Table 3.** Translational diffusion coefficient for D-and G-variants of SARS-CoV-2 (*D*_*v*_*10^15^/m^2^/s) due to heat energy only.

| T/k | $\eta/kg/m\ s$ | (D-variant) | (G-variant) |
| --- | --- | --- | --- |
| 293.15 | 1.801 | 2.398667407 | 2.183742183 |
| 298.15 | 1.600* | 2.7* | 2.5* |
| 303.15 | 1.433 | 3.065210576 | 2.838157940 |
| 308.15 | 1.292 | 3.455799930 | 3.199814750 |
| 313.15 | 1.173 | 3.868150269 | 3.581620619 |
| 318.15 | 1.066 | 4.324378068 | 4.004053766 |

\* The data for SARS-CoV-2 D and G variants are from the works of Al-kuraishy *et al.* (2022)

When particles of any size—macromolecules, supramolecules, and micro-molecules—interact with the cell membrane or with any other, two likely diffusivities are present. The first happens at the beginning of an attractive or repulsive interaction, mostly because of the influence of thermal energy. However, attractive interaction is of interest for the purpose of binding. The second is because the binding interaction slows down the molecule’s motion. Table 4 illustrates the increasing trend caused by rising temperature as stated earlier, even if values near binding are lower than those before binding at the start of the attractive interaction because of long-range forces. Compared to the G variant, the D variant of SARS-CoV-2 is faster.

**Table 4.** Translational diffusion coefficient (*D*_*v*_) for D-and G-variants of SARS-CoV-2 at the beginning of electrostatic interaction and upon close to binding at different temperatures (Table 3)

| The beginning |  | Close to binding |  |
| --- | --- | --- | --- |
| $D_{V(\epsilon_0\epsilon_r)}*10^{15}/\text{m}^2/\text{s}$ | | $D_{v(\epsilon_0)}*10^{15}/\text{m}^2/\text{s}$ | |
| D-variant | G-variant | D-variant | G-variant |
| 4.360 | 4.151 | 1.519 | 1.405 |
| 4.989 | 4.752 | 1.738 | 1.609 |
| 5.664 | 5.395 | 1.973 | 1.827 |
| 6.386 | 6.082 | 2.225 | 2.059 |
| 7.147 | 6.808 | 2.490 | 2.305 |
| 7.990 | 7.611 | 2.784 | 2.577 |

D- and G-variants of SARS-CoV-2 are those studied by Al-kuraishy et al. (2022); this also includes reference to viscosity coefficient (1.6 Pa.s) of nasal mucus in the literature (Leal et al., 2017).

Entry into host cell marks the first step of viral infection. The S protein first undergoes proteolysis into S1 and S2 (Heald-Sargent et al., 2012). This may be mediated either by host cell furin, by serine proteases such as the transmembrane protease, serine 2 (TMPRSS2) (Cheng et al., 2020), or by cathepsin proteases in the late endosome/endolysosomes (Peacock et al., 2021).

But regardless of the pathway—early, late, or delayed—the coronavirus’s inability to attach to the host cell membrane for whatever reason is a surefire means to disinfect the cell. The ultimate objective must always be to determine how to prevent or postpone the effective viral-membrane interaction that occurs before any type of binding. If such binding occurred, it must be made unstable, making it impossible for the coronavirus–membrane complex to be thermodynamically stable; it is therefore clear that Gibbs free energy and nonequilibrium binding energy are different parameters. A proper understanding of nonequilibrium binding energy and its formation could be helpful in designing a means of circumventing it.

Two aspects of the non-state function are examined based on the following premises: the first is the nonequilibrium binding energy, which starts at a point in space and time where the pathogens in general (but specifically SARS-CoV-2 for the purposes of this study) and the host cell membrane begin to interact attractively; the second is the point at which a vacuum-like space forms between the invading pathogen and the susceptible host cell as a result of the displacement of aqueous solvent from the cell membrane surface into the extracellular fluid.

Recall that in strictly thermodynamic terms, a “non-state function,” otherwise known as a “path function,” is a property that depends on the path or process taken to reach a particular state, unlike state functions, which only depend on the initial and final states, even if, as stated, one may be interested only in what transpired at the beginning and at the end when binding with higher stability has occurred, similar to the view elsewhere (Kao *et al*., 1993). The findings of this study demonstrate that the nonequilibrium binding energy (the lowest and maximum) for the D- and G-variants of SARS-CoV-2 is several times more than the thermal energy denoted by *R*_*g*_*T*, where *T* is between 293.15 and 318.15 K and *R*_*g*_ is the gas constant (Table 5). The omicron versions showed similar findings (Table 6).

**Table 5.** Minimum and maximum nonequilibrium binding energy (NEBE) of D- and G- variants of SARS-CoV-2.

| Min NEBE /cal/mol. |  | Max NEBE /kcal/mol. |  |
| --- | --- | --- | --- |
| D-variant | G-variant | D-variant | G-variant |
| 891.284 | 891.284 | 833.520 | 951.628 |
| 906.486 | 906.486 | 810.868 | 879.437 |
| 921.687 | 921.687 | 715.711 | 817.126 |
| 936.889 | 936.889 | 667.957 | 762.605 |
| 952.091 | 952.091 | 626.286 | 715.029 |
| 967.292 | 967.293 | 587.596 | 670.856 |

Min and Max stand for minimum and maximum respectively.

**Table 6.** Minimum and maximum nonequilibrium binding energy of Omicron & delta variants of SARS-CoV-2.

| Min NEBE /cal/mol. |  | Max NEBE /kcal/mol. |  |
| --- | --- | --- | --- |
| Omicron-variant | Delta-variant | Omicron-variant | Delta-variant |
| 906.486 | 906.486 | 732.584 | 603.776 |

Min and Max stand for minimum and maximum respectively. The $R_0$ , $R_{int}$ , and $C_v$ values are 53.03300859 nm, 75 nm, and 7.542087542 exp. (+20) respectively for delta variant; they are 64.34671709 nm, 91 nm, and 4.222320368 exp. (+20) respectively for omicron variant. $2\hat{R}_v$ (diameter) for delta variant is 75 nm (Vieira, *et al.*, 2024); diameter for omicron as a representative average for all variants is 91 nm (Ke *et al.*, 2020).

There is a need to opine that the lack of definite values for the molar masses and the diameters of coronaviruses of the kind explored in this study is a major challenge. For instance, Delta and Gamma variants of SARS-CoV-2 have, respectively, 75 nm and 79 nm diameters (Vieira *et al*., 2024); the average SARS-CoV-2 size is 91 +/- 11 nm (Ke *et al*., 2020).

These independent parameters have roles in the computation of dependent variables such as those indicated by Eqs (13), (17), etc. It is the nonequilibrium binding energy that is ultimately affected.

The respiratory tract is a key part of an elaborated line of defense based on a unique cellular ecosystem (Deprez *et al*., 2020). In this regard, the mucus in whatever location in the respiratory system plays a key role. Thus, secretory and multiciliated cells form a self-clearing mechanism that efficiently removes inhaled particles from the upper airways, impeding their transfer to deeper lung zones (Deprez *et al*., 2020). This can be overwhelmed with time. Although the nose and bronchus share many cellular properties, which has led to the definition of a pathophysiological continuum in allergic respiratory diseases (McDougall *et al*., 2008 & Samitas *et al*, 2018), they differ by features such as host defense against viruses, oxidative stress (Mihaylova *et al*., 2018), or antibacterial mechanisms (Roberts *et al*., 2018 & Giovannini-Chami *et al*., 2018). Be it as it may, individuals who are heavily infested with or without symptoms are liable to releasing viral particles in viral copies/m^3^ or viral copies/hr to the surrounding. In order to determine adequate viral number density after a long time exposure to the environment, this study made an effort for the first time to link breathing rate and breath emission to the question of coronavirus infection preceded by the binding interaction of viral particles with the membrane.

Exploring the breath emission in viral number density (number of viral particles per cubic meter), the multiplicity of infection (MOI i.e. *m* for convenience) with which initial number of infected cells (*N*_*IC*_) was computed, giving values as in Table 7. The initial viral number density (*C*_*v(τ*→0)_) in its dilute state was then determined and displaced in Table 7. The maximum (*R*_*int*(*τ*→0)_) and minimum (*R*_0(*τ*→0)_) inter viral-membrane distance under dilute condition and the same parameters under higher number density are also shown in Table 7. With the aforementioned parameters, the minimum (influenced by the relative permittivity (ε_*r*_) of the medium) and the maximum (free from ε_*r*_) nonequilibrium binding energies (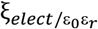 and 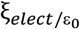) were computed for breath emission (ℬ) of 1.29 and 27.9 exp. (7) per cubic meter (Zheng, *et al*. 2022). The delta variant has greater values of NEBE than omicron variant.

**Table 7.**
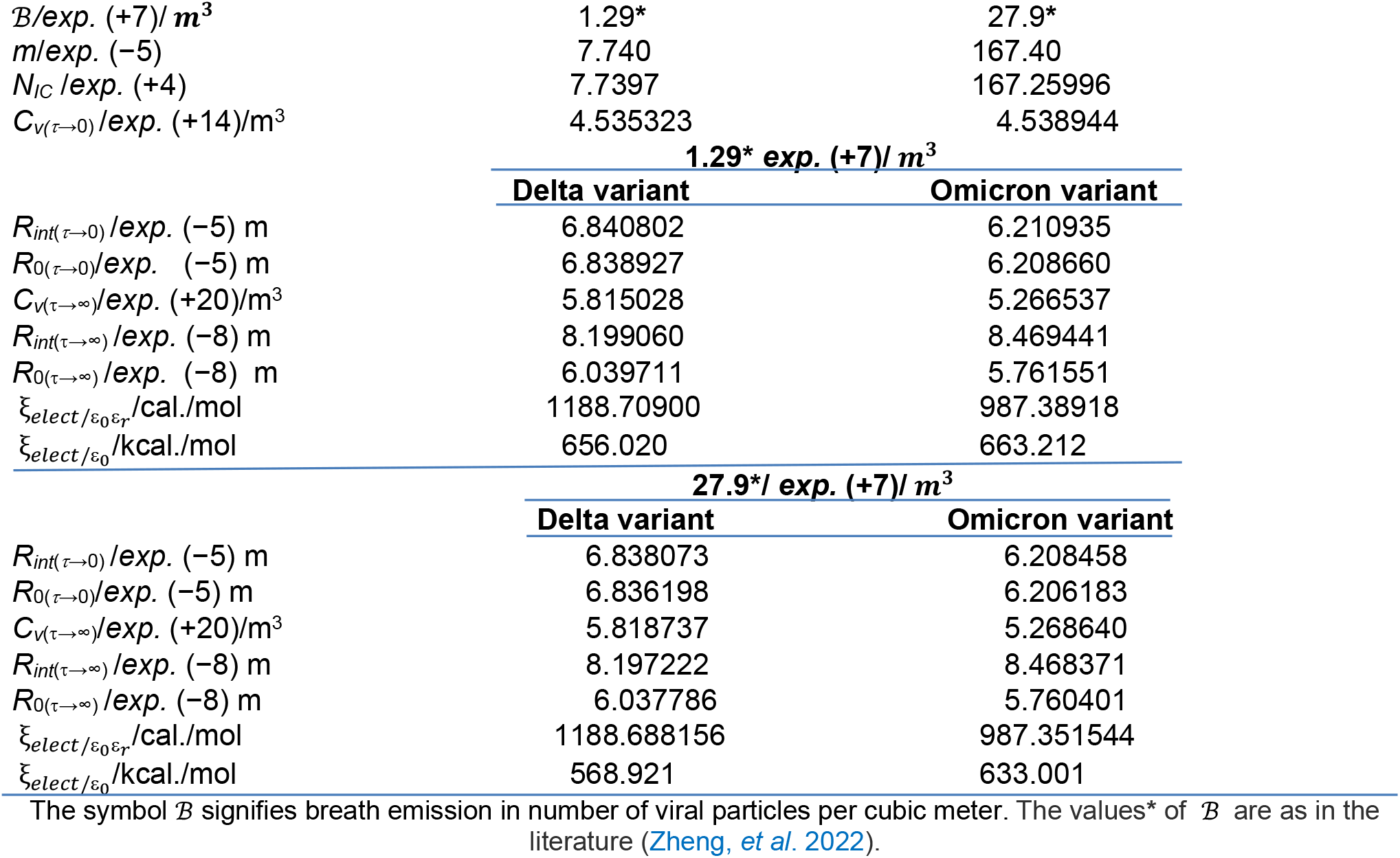
Physical parameters and cognate nonequilibrium binding energies of delta and omicron variants of SARS-CoV-2 given viral emission in viral copies per unit volume

Table 8 provides all physicochemical parameters for all computations based on breath emission rates in virus particles per hour, which vary from 9.31 to 201 exp. (6) /hr (Zheng et al. 2022). Results comparing the nonequilibrium binding energy of Delta and Omicron versions of SARS-CoV-2 are shown in Table 7, and Table 8a displays findings for the Omicron variant that are comparable to those in Table 7. These enables indirect comparison with literature values as follows: Omicron has the highest binding affinity at −128.35±10.91 kcal/mol, followed by Delta at −82.78±8.84 kcal/mol and the wild type at −73.26±7.46 kcal/mol, according to the results of a simulation (Ju et al., 2024). As long as the values correspond to the Gibbs free energy of binding to ACE2, this study’s findings support the idea that the Omicron variant has a lower energy barrier for binding than other variants due to its higher nonequilibrium binding energy (Table 7 & 8b), which is kinetic in nature and related to activation energy. This means that the omicron variant is likely to be more stable than other variants in its binding to the cell membrane.

**Table 8a.**
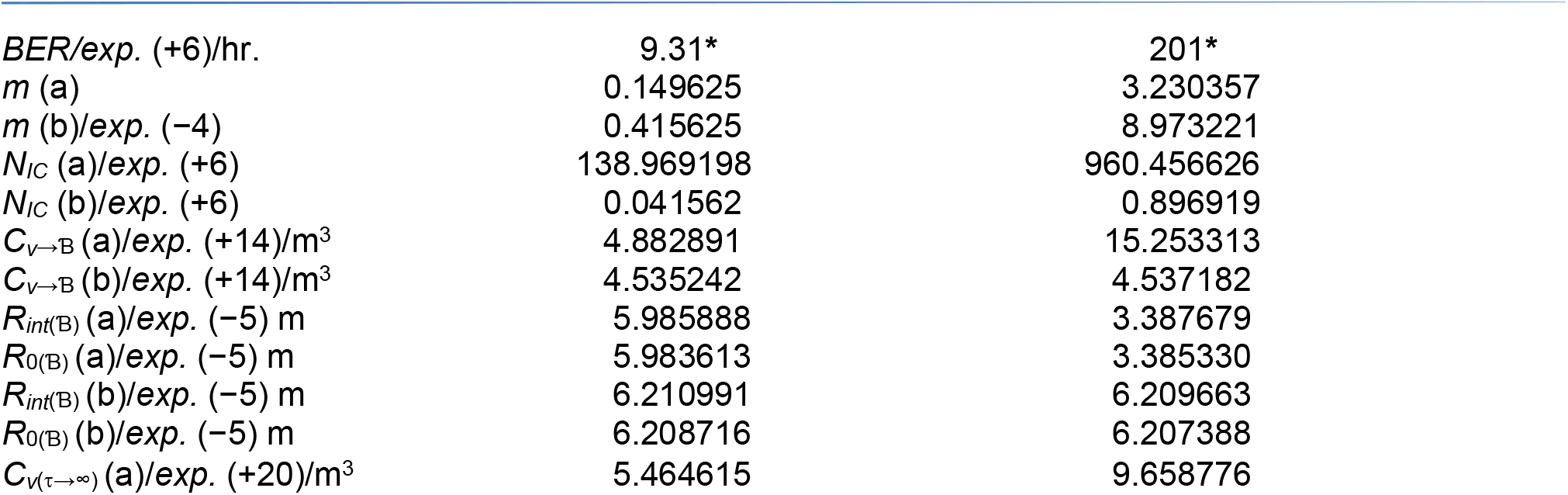
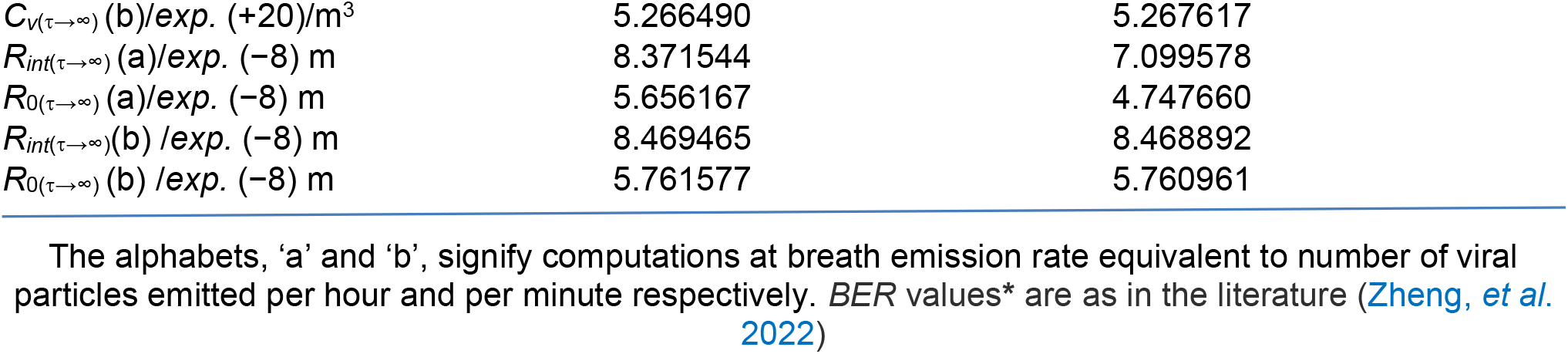
Physical parameters and cognate nonequilibrium binding energies of omicron variant of SARS-CoV-2 given breathe emission rate per hour.

**Table 8b.** Minimum and maximum nonequilibrium energies.

| Parameters | 9.31 exp. (6)/hr. | 201 exp. (6)/hr. |
| --- | --- | --- |
| $\xi_{elect/\varepsilon_0\varepsilon_r}(a)/cal./mol$ | 522.390979 | 962.988964 |
| $\xi_{elect/\varepsilon_0}(a)/kcal./mol$ | 651.703 | 444.157 |
| $\xi_{elect/\varepsilon_0\varepsilon_r}(b)/cal./mol$ | 987.188601 | 987.273963 |
| $\xi_{elect/\varepsilon_0}(b)/kcal/mol$ | 683.353 | 663.212 |

The alphabets, 'a' and 'b' are as previously defined.

It is necessary to comment that different approaches give different Gibbs free binding energies going by the following: Using unfamiliar method, involving a homogeneous glycan setup, the binding free energy (Δ*G*_bind_) for Omicron was −30.21±5.48 kcal/mol lower than −18.32±1.62 kcal/mol for the WT (Nguyen et al., 2022); this inference based on number theory needs to be seen as one in which higher negative magnitude of the mixed decimal signifies greater thermodynamic stability of feasibility of ligand-receptor complex. Besides they are much lower than Ju et al. (2024) report. The binding of SARS-CoV-RBD and SARS-CoV-2-RBD to ACE2-PD with time, is driven by electrostatic interactions (Nguyen *et al*., 2020). This is significant because it reminds researchers in science and medicine that it is possible that there may be other existing variants with stronger binding affinity that should thus, require stronger drugs or medications that can hinder binding processes. These findings are based on coarse-grained and all-atom steered (MD) simulations experiments which showed that SARS-CoV-2 (− 875.89 kcal/mol.) had greater binding affinity to its receptor than SARS-CoV (− 695.95 kcal/mol.). It is not certain whether those values reported as electrostatic energies are free energies for binding. If they are total energies according to convention, then the kinetic components should be equal in magnitude to the former but opposite in sign. In this way, they compare very well with the results in this study. Some examples are as follows: 670.6–951.6 and 587.6–833.5 kJ/mol for G and D variants of SARS-CoV-2, respectively (Table 5).

However, according to the Nudged Elastic Band (NEB) binding energy profiles, Delta has the lowest transition state barrier at 9.21 kcal /mol, whereas the wild type has the highest at 17.87 kcal /mol. Although the Omicron form is said to have a little greater energy barrier than Delta variants, no precise figure was provided (Ju et al., 2024). These values are completely out of line with nonequilibrium binding energies like 632.7–719.9 kcal/mol (Table 7), which span the Delta and Omicron variants and indicate the amount of work required to overcome mechanical kinetic energy in order to hold the ligand to the receptor.

### 4.1 Viscosity as a measure of resistance of the medium on mobility of biomolecules and its variation

The nature of the medium should not be the only factor considered when evaluating viscosity as a measure of resistance to any particle motion in the medium; a medium that produces high viscosity for one solute may do so at a lower viscosity for another solute with distinct physicochemical properties. In contrast to solely polar solutes lacking lipophilic functional groups, polar solutes with lipophilic properties can diffuse more quickly through lipid bilayers that contain peripheral and integral proteins, glycosylated and unglycosylated proteins, etc. It is anticipated that such a solvent or solute will have a higher resistance than the former. Water (kinetic size: 2.68Å) and ethanol (kinetic size: 4.5Å) have diffusivities over the chitosan membrane of 2.8 and 3.32 exp. (−11) m^2^/s, respectively (Dudek & Borys, 2019) which correspond to viscosities of 0.0303 and 0.01524 kg/m/s calculated in this study. The conditions in the cytoplasm and mucus are pertinent even if this is about the membrane. The scientific and medical (therapeutic) implications are that one SARS-CoV-2 variant diffuses more quickly than the other, making it more likely to strike its target membrane and, if binding is stable, more contagious. Therefore, such a situation necessitates the use of more potent, highly effective medications as well as alternative therapeutic approaches.

Viscosity has a major impact on the velocity of translational diffusion in cells, which in turn affects cellular reactivity by influencing the frequency and manner of reactant collisions with enzyme active sites (O’Loughlin et al., 1980; Kao et al., 1993). Diffusion coefficients for macromolecules (Wojcieszyn et al., 1981) and tiny solutes (Kao et al., 1993; O’Loughlin et al., 1980) are 5–50 times lower in mammalian cells than in pure water, according to research. Meanwhile, the mucus consists of two layers, with the upper layer being a gel with a 3% mucin network and the remaining 90-95% water with electrolytes, serum proteins, immunoglobulins, and lipids (Verdugo, 1990). However, experimental studies show mucus viscosity can be as high as 10,000 times that of water (Mestecky et al., 2005) due to overlapping adhesive mucin fibers. It seems unimaginable that the diffusivity of SARS-CoV-2 in mucus could be similar to that in water, and in particular, if normal body temperature remains consistent without variation. As initially cited by Adamczyk et al. (2021), this and the study’s results (Table 3) also support the theory that the virion diffusion coefficient is substantially lower in media with a higher viscosity, like mucus or saliva, which adds to the overall virion transfer rate (Roselli & Diller, 2011). The cellular domains containing dimer 1 (a fluorescent molecular rotor and an effective PDT photosensitizer) had a viscosity of (300±50) cP after photodynamic treatment (PDT). This number is higher than the 50 cP value that was achieved before PDT (Kuimova, 2008/2009). The regulation of mucus and cytoplasmic viscosity may benefit from this discovery. Viral, medication, and Ig diffusivities can be controlled by increasing or decreasing the mucus’s viscosity.

### 4.2 General matters arising

The nonequilibrium binding energy is not just about the binding of a substrate and ligand to an enzyme and a receptor, respectively, but it also includes biological porters, i.e., carriers such as carrier proteins, albumin, lipid nanoparticles (LNP), etc. In order to deliver, these carriers must make contact with the target surface, usually the cell or receptor membrane. Examples of lipid-based nanoparticles as potential carriers of medications are liposomes, nanoemulsions, solid lipid nanoparticles (LNPs), nanostructured lipid carriers, and lipid polymer hybrid nanoparticles (Mehta et al., 2023). The importance and usefulness of LNPs lie in their capacity to safeguard drugs from *in vivo* degradation, boost their solubility and efficacy, enable targeted drug delivery to the disease site, regulate drug release, and alter drug bio-distribution (Shah, 2020). The most crucial element is targeted delivery, which cannot be accomplished by “saltatory mechanisms” or *American James Bond flyover action movies, frequently to the delight of teenagers*. In this case, the RMSD is crucial because it allows investigators to forecast the distance that free drugs or LNPs can travel in space before being completely distributed at equilibrium and outside of the equilibrium state as they approach the target.

In addition to the viscosity (either known or unknown) of the biological medium, there are other factors that may affect the upward trajectory of the rate of diffusion of the nanoparticles. The effectiveness of such nanoparticles in therapeutic applications is based on their size, shape, and composition, which are morphological characteristics in addition to the surface chemistry (net charge in particular) that slow down the transport of bioactive cargo, which adversely alters therapeutic efficacy (Ridolfo et al., 2021). Using Line-FRAP measurements, Brownian dynamics simulations, and molecular docking, researchers found that the diffusion rates of the small molecules are highly affected by self-aggregation, interactions with the proteins, and surface adsorption (Dey et al., 2022). Some drugs (e.g., quinacrine) have their diffusion unaffected by protein crowders (Dey et al., 2022). Based on these findings, it might be useful to create neutral carriers for medicines (like nanoparticles) that can move easily in mucus, while others could be designed to move less freely because of their interactions with mucus components, including the coronavirus.

Nonequilibrium binding energy benefits both mucosa and cytoplasm, with fluid-like and gel-like compartments. Mucus also has gel-like consistency, suggesting two intra-mucus compartments (*Ugwoke et al*., *2005, Mistry et al*., *2009, Gizurarson, 2015*). The effects of crowding arise from two phenomena, hard-core repulsions and nonspecific chemical (soft) interactions (Fodeke & Minton, 2011, Knowles et al., 2011, Miklos et al., 2011, Phillip & Schreiber, 2013, Sarkar & Pielak, 2013, Wang et al., 2012, Zhou, 2013.). Hard-core repulsions limit the spatial volume of biological macromolecules, favoring compact forms over expanded ones, resulting in increased viscosity. Additionally, the presence of crowding molecules not only reduces available volume or volume exclusion effects (Aumiller et al., 2014), but also engages in chemical interactions. Increasing medium viscosity slows viral particle diffusion, ensuring drugs, antibodies, and carriers (LNP) maintain their mobility and affinity for pathogens. However, attractive interactions may destabilize essential enzymes, potentially hindering viral replication, as seen with chymotrypsin inhibitor 2 (Wang et al., 2012, Minton, 2013) in reconstituted *E. coli* cytosol (Sarkar et al., 2013).

## 5.0 CONCLUSION

A more practical method for determining the viral number density in the order of exp. (+20) was calculated using the successfully derived equations for the multiplicity of infection (*MOI*) associated with breath emission (viral number per hour or per cubic meter) and ventilation rate. The size and molar mass of the virus need to be consistent to enable accurate comparison of nonequilibrium binding energy (NEBE) among variants since the latter is a function of those parameters. If there are no computation errors, the values of minimum NEBE 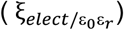 remain constant because temperature largely controls the minimum NEBE. As is typical, diffusivities increase with temperature; however, near binding, the binding limitations cause the diffusivities to be lower than when they are far from binding.

The decreasing trend in maximum NEBE 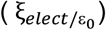 with rising temperature implies that the binding affinity of the virus may be attenuated at higher temperatures. It is, therefore, medically plausible to administer airborne, parenteral, oral, etc., drugs at temperatures above body temperatures that are tolerable and in a controlled fashion. As long as variants of SARS-CoV-2 exhibit different values of 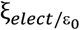, the logical response should be the continuation of searches for better drugs and vaccines.

Future studies must, however, clearly identify the method or processes for calculating those parameters as well as the molar masses of the coronavirus variants with specific values generated for them in order to compare their NEBEs with accuracy. This is due to the fact that various techniques produce varying viral sizes for the same variant and viscosities of a particular medium (cytoplasm, mucus, etc.).

## DISCLAIMER (ARTIFICIAL INTELLIGENCE)

Author(s) hereby declare that NO generative AI technologies such as Large Language Models (ChatGPT, COPILOT, etc.) and text-to-image generators have been used during the writing or editing of this manuscript.

## DEDICATION

The burning issue of equal opportunity for all without divisive and derisive policies was the main concern of Professor Ambrose Folorunsho Alli, the former governor of the now-defunct Bendel State, and his deputy, Chief Demas Onoliobakpovba Akpore. They made textbooks and even writing materials available for all regardless of social status; they made the state look like a one-party state not by manipulation or coercion, but by being statesmen for all in line with constitutional order; they separated the state from personality cult; hence, the governor never customized supplies, educational material, medical material, etc. after his name; rather, such supplies were labeled, “These are properties of the state and not for sale.” This is an expression of selfless service to humanity.

## ACKNOWLEDGEMENT

I am very grateful to my siblings for their financial and in-kind support. Grammar-checking services by QuillBot Company are also appreciated.

## AUTHORS’ CONTRIBUTIONS

The sole author designed, analyzed, interpreted and prepared the manuscript.

## CONSENT

NA

## ETHICAL APPROVAL (WHERE EVER APPLICABLE)

NA

